# A smoothed version of the Lassosum penalty for fitting integrated risk models

**DOI:** 10.1101/2021.03.09.434653

**Authors:** Georg Hahn, Dmitry Prokopenko, Sharon M. Lutz, Kristina Mullin, Rudolph E. Tanzi, Christoph Lange

**Affiliations:** Harvard T.H. Chan School of Public Health; Genetics and Aging Research Unit, McCance Center for Brain Health, Department of Neurology, Massachusetts General Hospital, Boston, MA 02114; Harvard University; Harvard School of Public Health

**Keywords:** Integrated risk model, Lassosum, Nesterov, Polygenic Risk Scores, Smoothing

## Abstract

Polygenic risk scores are a popular means to predict the disease risk or disease susceptibility of an individual based on its genotype information. When adding other important epidemiological covariates such as age or sex, we speak of an integrated risk model. Methodological advances for fitting more accurate integrated risk models are of immediate importance to improve the precision of risk prediction, thereby potentially identifying patients at high risk early on when they are still able to benefit from preventive steps/interventions targeted at increasing their odds of survival, or at reducing their chance of getting a disease in the first place. This article proposes a smoothed version of the “Lassosum” penalty used to fit polygenic risk scores and integrated risk models. The smoothing allows one to obtain explicit gradients everywhere for efficient minimization of the Lassosum objective function while guaranteeing bounds on the accuracy of the fit. An experimental section on both Alzheimer’s disease and COPD (chronic obstructive pulmonary disease) demonstrates the increased accuracy of the proposed smoothed Lassosum penalty compared to the original Lassosum algorithm, allowing it to draw equal with state-of-the-art methodology such as LDpred2 when evaluated via the AUC (area under the ROC curve) metric.

## 1 Introduction

Polygenic risk scores are a statistical aggregate of risks typically associated with a set of established DNA variants. If only genotype information of an individual is used to predict its risk, we speak of a polygenic risk score. A polygenic risk score with added epidemiological covariates (such as age or sex) is called an integrated risk model (Wand et al., 2020). The goal of both polygenic risk scores and integrated risk models is to predict the disease risk of an individual, that is the susceptibility to a certain disease. Such scores are usually calibrated on large genome-wide association studies (GWAS) via high-dimensional regression of a fixed set of genetic variants (and additional covariates in case of an integrated risk model) to the outcome. In this article, we focus on the more general case of an integrated risk model.

Since the potential for broad-scale clinical use to identify people at high risk for certain diseases has been demonstrated (Khera et al., 2018), polygenic risk scores and integrated risk models have become a widespread tool for the early identification of patients who are at high risk for a certain disease and who could benefit from intervention measures. However, the accuracy of current polygenic risk scores, measured with the AUC metric (Area under the ROC Curve, where ROC stands for receiver operating characteristic, see Mandrekar (2010)), varies substantially for important diseases. For instance, the AUC achieved by state-of-the-art methods ranges from around 0.8 for type 1 diabetes to around 0.7 for coronary artery disease and schizophrenia (Mak et al., 2017), while for atrial fibrillation the AUC is around 0.64 (Huang and Darbar, 2017), a value which is considered less than acceptable (Mandrekar, 2010; Hosmer and Lemeshow, 2000). For this reason, increasing the accuracy of scores is desirable, which is the focus of the proposed smoothing approach.

Several methodological approaches have been considered in the literature to compute a polygenic risk score or an integrated risk model for a given population, and to predict a given outcome (disease status). For instance, LDpred of Vilhjálmsson et al. (2015) and LDpred2 of Privé et al. (2019) fit a Bayesian model to the effect sizes via Gibbs sampling, and obtain a score via posterior means of the fitted model. The PRS-CS approach of Ge et al. (2020) likewise utilizes a high-dimensional Bayesian regression framework in connection with a continuous shrinkage prior (hence the suffix *CS* for continuous shrinkage) on SNP effect sizes. Fitting genotype data to a disease outcome can also be achieved by means of a simple penalized regression using the least absolute shrinkage and selection operator (Lasso) of Tibshirani (1996), for instance using the *glmnet* package on CRAN, see Friedman et al. (2010, 2020).

One popular way to fit a polygenic risk score is the “Lassosum” approach of Mak et al. (2017). Note that in Mak et al. (2017), no integrated risk models are considered. The Lassosum method is based on a reformulation of the linear regression problem ***y*** = *X****β*** + ***ϵ***, where *X* ∈ ℝ^*n*×*p*^ denotes SNP data for *n* individuals and *p* SNP locations, ***y*** ∈ ℝ^*n*^ denotes a vector of outcomes, ***β*** ∈ ℝ^*p*^ is unknown, and ***ϵ*** ∼ *N*_*n*_(0, *σ*^2^*I*_*n*_) is an *n*-dimensional, independently and normally distributed error term with mean zero and some variance *σ*^2^ *>* 0 (where *I*_*n*_ denotes the *n*-dimensional identity matrix). The authors start with the classic Lasso objective function *L*(***β***) = /||***y*** − *X****β***/||_2_ + 2*λ*/||***β***/||_1_, where *λ* ≥ 0 denotes the Lasso regularization parameter controlling the sparseness of the solution, and rewrite it using the SNP-wise correlation ***r*** = *X*^T^***y*** as

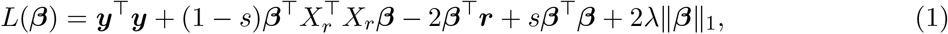

where *X*_*r*_ denotes the matrix of genotype data used to derive estimates of LD (linkage disequilibrium), *λ* ≥ 0 is the Lasso regularization parameter controlling the sparseness of the estimate, and *s* ∈ (0, 1) is an additional regularization parameter used to ensure stability and uniqueness of the Lasso solution. Importantly, in Mak et al. (2017) the authors derive an iterative scheme to carry out the minimization of eq. (1) which only requires one column of *X*_*r*_ at a time, thus avoiding the costly computation of the matrix *X*_*r*_^T^*X*_*r*_ ∈ ℝ^*p*×*p*^.

In this work, we consider a different approach for minimizing eq. (1). Using the methodology of Nesterov (2005), we propose to smooth the non-differentiable *L*_1_ penalty in eq. (1), thus allowing us to compute explicit gradients of eq. (1) everywhere. This in turn allows us to efficiently minimize the Lassosum objective function using a quasi-Newton minimization algorithm such as BFGS (Broyden–Fletcher–Goldfarb–Shanno). Besides enabling a more efficient and more accurate computation of the score, our work extends the one of Mak et al. (2017) in that we do not solely consider polygenic risk scores, but the more general integrated risk models. Our approach follows as a special case from Hahn et al. (2020b,a), who propose a general framework to smooth *L*_1_ penalties in a linear regression. Importantly, employing a smoothing approach has a variety of theoretical advantages following directly from Hahn et al. (2020b). Apart from obtaining explicit gradients for fast and efficient minimization, the smoothed objective is convex, thus ensuring efficient minimization, and it is guaranteed that the solution (the fitted integrated risk model) obtained by solving the smoothed Lassosum objective is never further away than a user-specified quantity from the original (unsmoothed) objective of Mak et al. (2017).

We evaluate all aforementioned approaches by computing an integrated risk model in two experimental studies, one on Alzheimer’s disease using the summary statistics of Kunkle et al. (2019) and Jansen et al. (2019), and on COPD (chronic obstructive pulmonary disease) using FEV1 data of Regan E.A. (2010); NHLBI TOPMed (2018). In the first case, the response is binary, whereas in the second study the response is continuous. Our simulations demonstrate that smoothing the Lassosum objective function results in a considerably enhanced performance of the Lassosum approach, allowing it to draw equal with approaches such as LDpred2 or PRScs.

Analogously to the original Lasso of Tibshirani (1996), the *L*_1_ penalty employed in eq. (1) causes some entries of arg min_*β*∈ℝ*p*_ *L*(***β***) to be shrunk to zero exactly (provided the regularization parameter *λ* is not too small). Therefore, Lassosum performs fitting of the polygenic risk score or integrated risk model and variable selection simultaneously.

This article is structured as follows. Section 2 introduces the smoothed Lassosum objective function and discusses its minimization, the theoretical guarantees it comes with, and its drawbacks. Section 3 evaluates the proposed approach, the original Lassosum approach, as well as additional state-of-the-art methods in two experimental studies on both Alzheimer’s disease and COPD. The article concludes with a discussion in Section 4. The appendix contains two figures showing plots of principal components for the genotype dataset employed in Section 3.1.

The methodology of this article is implemented in the R package *smoothedLasso* (see function *prsLasso* in the package), available on CRAN (Hahn et al., 2020c).

## 2 Methodology

The Lassosum function of eq. (1) consists of a smooth part, given by ***y***^T^***y*** + (1 − *s*)***β***^T^*X*_*r*_^T^*X*_*r*_***β*** − 2***β***^T^***r*** + *s****β***^T^***β***, and a non-smooth part, the *L*_1_ penalty 2*λ*/||***β***/||_1_. Only the latter needs smoothing, which we achieve with the help of Nesterov smoothing introduced in Section 2.1. Section 2.2 applies the Nesterov methodology to Lassosum and introduces our proposed smoothed Lassosum objective function. The proposed smoothed Lassosum actually follows from the more general framework of Hahn et al. (2020a,b). We demonstrate this in Section 2.3, where we also state the theoretical guarantees following from the framework.

### 2.1 Brief overview of Nesterov smoothing

In Nesterov (2005), the author introduces a framework to smooth a piecewise affine and convex function *f* : R^*q*^ → R, where *q* ∈ N. Since *f* is piecewise affine, it can be written for ***z*** ∈ ℝ^*q*^ as

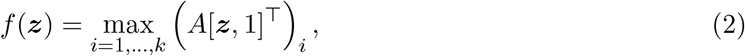

using *k* ∈ N linear pieces (components), where [***z***, 1] ∈ ℝ^*q*+1^ denotes the vector obtained by concatenating ***z*** and the scalar 1. In eq. (2), the linear coefficients of each of the *k* linear pieces are summarized as a matrix *A* ∈ ℝ^*k*×(*q*+1)^ (with the constant coefficients being in column *q* + 1).

The author then introduces a smoothed version of eq. (2) as

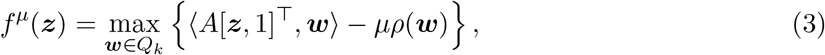

where 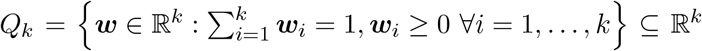 is the unit simplex in *k* dimensions. The parameter *µ* ≥ 0 controls the smoothness of the approximation *f*^*µ*^ to *f*, called the Nesterov smoothing parameter. Larger values of *µ* result in a stronger smoothing effect, while the choice *µ* = 0 recovers *f* ^0^ = *f*. The function *ρ* is called the proximity function (or prox-function) which is assumed to be nonnegative, continuously differentiable, and strongly convex.

Importantly, *f*^*µ*^ is both smooth for any *µ >* 0 and uniformly close to *f*, that is the approximation error is uniformly bounded as

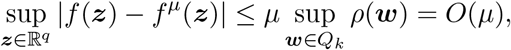

see (Nesterov, 2005, Theorem 1). Though several choice of the prox-function *ρ* are considered in Nesterov (2005), we fix one particular choice (called the entropy prox-function) in the remainder of the article for the following reasons: (a) The different prox-functions are equivalent in that all choices yield the same theoretical guarantee and performance; and (b) the entropy prox-function leads to a closed-form expression of eq. (3) given by

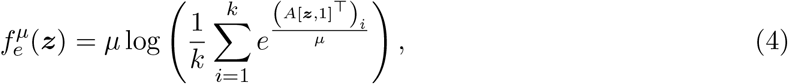

which satisfies the uniform bound

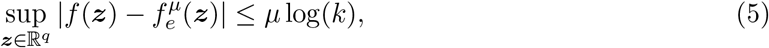

see Nesterov (2005) and Hahn et al. (2020a,b).

### 2.2 A smoothed version of the Lassosum objective function

As observed at the beginning of Section 2, it suffices to smooth the non-differentiable penalty 2*λ*‖***β***‖_1_ of the Lassosum objective function, where 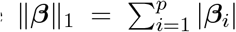 To this end, we apply Nesterov smoothing to each absolute value independently.

We observe that the absolute value can be expressed as piecewise affine function with *k* = 2 components, given by *f* (*z*) = max{−*z, z*} = max_*i*=1,2_ (*A*[*z*, 1]^T^)_*i*_, where

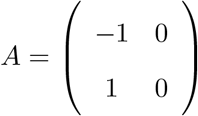

and *z* ∈ ℝ is a scalar. Substituting this specific choice of *A* into eq. (4) leads to a smoothed approximation of the absolute value given by

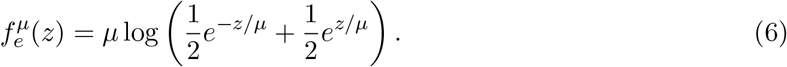

Substituting the absolute value in the *L*_1_ norm in eq. (1) with the approximation in eq. (6) results in a *smoothed version of the Lassosum objective function*, given by

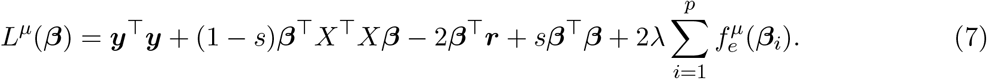

The first derivative of 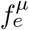 is explicitly given by

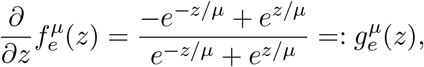

see also Hahn et al. (2020a,b), from which the closed-form gradient of the smoothed Lassosum objective function of eq. (7) immediately follows as

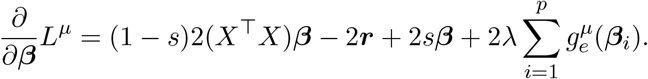

Using the smoothed version of the Lassosum objective function, given by*L*^*µ*^, and its explicit gradient 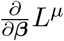 an integrated risk model can easily be computed by minimizing *L*^*µ*^ using a quasi-Newton method such as BFGS (Broyden–Fletcher–Goldfarb–Shanno), implemented in the funcition *optim* in *R* (R Core Team, 2014).

In eq. (7), the quantity *X* is not limited to contain only genotype information. Any data on the individuals (including additional epidemiological covariates) to compute the integrated risk model can be summarized in *X*. The other quantities in eq. (7) are the outcome ***y*** (either binary/discrete or continuous), the correlations ***r*** = *X*^T^ ***y***, and the additional regularization parameter *s* ∈ (0, 1) introduced by Mak et al. (2017) used to ensure stability and uniqueness of the Lasso solution.

### 2.3 Theoretical guarantees

Using the fact that the absolute value can be expressed as a piecewise affine function with *k* = 2, see Section 2.2, the error bound of eq. (5) can be re-written as

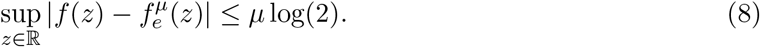

Since in our proposed smoothed version of eq. (7), only the non-smooth *L*_1_ contribution of the original Lassosum objective function of eq. (1) has been replaced, the bound of eq. (8) immediately carries over to a bound on the smoothed Lassosum. In particular,

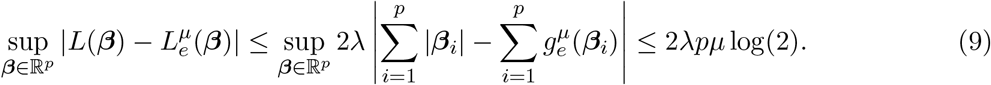

For a given computation of an integrated risk model, the Lasso parameter *λ >* 0 and the dimension *p* are fixed by the problem specification. According to eq. (9), this allows one to make the approximation error of our proposed smoothed Lassosum to the original Lassosum arbitrarily small as the smoothing parameter *µ* → 0.

As stated in Section 2.1 of Mak et al. (2017), the Lassosum objective of eq. (1) is equivalent to a Lasso problem, in particular its convexity is preserved. According to Proposition 2 in Hahn et al. (2020b), the smooth approximation of eq. (7) obtained via Nesterov smoothing is strictly convex. Since strictly convex functions have one unique minimum, and since a closed-form gradient 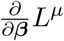 of *L*^*µ*^ is available (see Section 2.2), this makes the minimization of our proposed smoothed Lassosum in lieu of the original Lassosum very appealing.

Furthermore, two additional properties of eq. (7) can be derived from (Hahn et al., 2020b, Section 4.3). First, the arg min_*β*∈ℝ*p*_ *L*^*µ*^(***β***) is continuous with respect to the supremum norm (Hahn et al., 2020b, Proposition 4), which implies that the minimum of our proposed smoothed Lassosum *L*^*µ*^ converges to the one of the original Lassosum as *µ* → 0. Second, in addition to this qualitative statement, the error between the minimizers of the smoothed and original Lassosum function can be quantified a priori (Hahn et al., 2020b, Proposition 5).

## 3 Experiments

The proposed smoothed Lassosum approach is obtained by applying Nesterov smoothing to the *L*_1_ penalty of the Lassosum objective function, see eq. (1). A detailed study on the behavior of Nesterov smoothing applied to an *L*_1_ penalty using synthetic data can be found in Hahn et al. (2020b).

In this section, we evaluate the the performance of our proposed smoothed Lassosum approach of Section 2.2 in two experimental studies, one fitting an integrated risk model to binary outcomes in the context of Alzheimer’s disease (Section 3.1), and one fitting an integrated risk model to continuous outcomes in the context of COPD (Section 3.2). We benchmark our smoothed Lassosum approach, which we refer to as “SmoothedLassosum”, against the following state-of-the-art approaches:

1. We benchmark against the original Lassosum of Mak et al. (2017), implemented in the R package *lassosum* (Mak et al., 2020), and refer to it as “Lassosum”.
2. LDpred and LDpred2 (Vilhjálmsson et al., 2015; Privé et al., 2019) compute a polygenic risk score (but not integrated risk model) by inferring the posterior mean effect size of each marker by using a prior on effect sizes and LD information from an external reference panel. To run LDpred2, we employed the implementation in the R package bigsnpr on CRAN (Privé et al., 2020) using the Gibbs sampler to estimate effect sizes. We will refer to this algorithm as “LDpred2”.
3. PRScs of Ge et al. (2019) utilizes a high-dimensional Bayesian regression framework which places a continuous shrinkage prior on SNP effect sizes, an innovation which the authors claim is robust to varying genetic architectures. We use the implementation of Ge et al. (2020) and refer to it as “PRScs”.
4. We employ a simple penalized regression using the Lasso of Tibshirani (1996) to fit the genotype data to disease outcome. We employ the *Glmnet* package on CRAN, see Friedman et al. (2010, 2020). We will refer to this method as “Glmnet”.
5. We employ the unsmoothed Lasso of Hahn et al. (2020a), implemented in the R package *smoothedLasso* on CRAN (Hahn et al., 2020d). We refer to this method as “Lasso”.
6. Similarly, we also employ the smoothed Lasso of Hahn et al. (2020a), which is likewise implemented in *smoothedLasso* on CRAN (Hahn et al., 2020d). We refer to this method as “SmoothedLasso”.
7. We train a neural network with the *Keras* interface (Falbel et al., 2020a) to the *Tensorflow* machine learning platform (Falbel et al., 2020b). We train a network with four layers, having 20, 8, 4 and 2 nodes. We employ the LeakyReLU activation function, a dropout rate of 0.1, a validation splitting rate of 0.1, the *he normal* truncated normal distribution for kernel initialization, and kernel, bias and activity regularization with *L*_1_ penalty. The last layer employs the sigmoid (for Section 3.1) or ReLU (for Section 3.2) activation functions. The model is compiled for binary crossentropy loss (for Section 3.1) or mean absolute error loss (for Section 3.2) using the Adam optimizer, evaluated with the AUC (for Section 3.1) or the mean squared error (for Section 3.2) using 1000 epochs. We refer to the neural network as “NeuralNetwork”.
8. We employ *SBayesR* of Lloyd-Jones et al. (2019), a linear regression likelihood which takes into account GWAS summary statistics and a reference LD correlation matrix, and is coupled to a finite mixture of normal priors on the genetic effects. The normal priors allow one to incorporate sparsity and to perform Bayesian posterior inference on the model parameters, such as genetic effects, variance components and mixing proportions. The method is implemented in the toolbox *GCTB* (Zeng et al., 2020). *We employ SBayesR with default parameters and refer to it as “SBayesR”*.
9. *We run MegaPRS* of Zhang et al. (2021). In particular, we employ the robust version *Bolt Predict*, as suggested by the authors, using default parameters given the example section of MegaPRS (a cross validation proportion of 0.1, the *--ignore-weights* option and a power parameter of −0.25). MegaPRS is implemented in the *LDAK* package (Speed, 2021). We refer to it as “MegaPRS”.
10. We use epidemiological covariates only in a simple linear regression fit to the response. We refer to this as “EpiOnly”.

The Lassosum, LDpred2, PRScs, SBayesR, and MegaPRS algorithms are only designed to fit polygenic risk scores, but not integrated risk models. To include epidemiological covariates for these methods (and thus fit an integrated risk model), we first perform a linear regression of the epidemiological covariates to the outcome, and then run the aforementioned methods on the residuals. Importantly, in order to apply Lassosum with epidemiological covariates, we additionally have to recompute the SNP-wise correlation ***r*** = *X*^T^ ***y*** as in eq. (1) using the residuals in place of ***y***.

Note that Glmnet, as well as Lasso and SmoothedLasso, can be applied in two ways. First, they can be applied to both the epidemiological covariates and genotype information in one go, given all information is summarized in the design matrix. Second, they can likewise be applied to residuals after regressing out all epidemiological covariates. For consistency with the way the Lassosum, LDpred2, PRScs, SBayesR, and MegaPRS algorithms are applied, we also employ Glmnet, Lasso and SmoothedLasso to residuals after regressing out all epidemiological covariates. Throughout the section, we fix the Lasso regularization parameter at *λ* = 2^-3^, the Lassosum regularization parameter *s* in eq. (1) at *s* = 0.5 (this parameter is used to ensure stability and uniqueness of solution), and the smoothing parameter of Section 2.2 at *µ* = 0.1.

### 3.1 Alzheimer study

We performed training and testing of different PRS algorithms using summary statistics for Alzheimer’s disease (AD), together with genotype data imputed on the Haplotype Reference Consortium (HRC), see McCarthy et al. (2016). The HRC-imputed genotype data was downloaded from Partners Biobank (Partners, 2020) (described below). The summary statistics are matched to genotype data for chromosomes 1–22 of 2, 465 patients available in the Partners Biobank. As initial training weights we considered two sets of summary statistics from two largest available AD GWAS: the one of clinically defined AD cases of Kunkle et al. (2019), and the one of AD-by-proxy phenotypes of Jansen et al. (2019).

The dataset of Kunkle et al. (2019) contains a total of 11, 480, 632 summary statistics, given by p-value, effect size (beta), and standard deviation of the effect size. Each entry is characterized by its chromosome number, position on the chromosome, as well as the effect allele and non-effect allele. The dataset of Jansen et al. (2019) contains a total of 13, 367, 299 summary statistics in the same format as the one of Kunkle et al. (2019).

Partners Biobank is a hospital-based cohort from the MassGeneral Brigham (MGB) hospitals. This cohort includes collected DNA from consented subjects linked to electronic health records. We have obtained a subset in April 2019, which included AD cases and controls. Cases were defined as subjects who were diagnosed with AD based on the International Statistical Classification of Diseases and Related Health Problems (ICD-10), see World Health Organization (2021). Controls were selected as individuals of age 60 and greater, who had no family history of AD, no diagnosed disease of nervous system (coded as G00-G99 in ICD-10), no mental and behavioral disorders (coded as F01-F99 in ICD-10), and a Charlson Age-Comorbidity Index of 2, 3, or 4 (Charlson et al., 1994; Karlson et al., 2016).

We performed the following quality control steps on the HRC-imputed genotype data from Partners Biobank. Relatedness was assessed with KING (Manichaikul et al., 2010; Chen, 2021) and population structure was assessed with principal components. Principal components were calculated on a pruned subset (PLINK2 parameters: --indep-pairwise 50 5 0.05) of common variants (MAF *>* 0.1). We excluded subjects which had a KING kinship coefficient *>* 0.0438 (third degree of relatedness or less) and which were at least 5 standard deviations away from the mean value of the inbreeding coefficient. We kept only self-reported non-hispanic white (NHW) individuals and excluded outliers, defined as subjects which are at least 5 standard deviations away from the mean value of each of the ten principal components (see Section A). There was a total of 2, 465 subjects (481 cases) left for analysis.

To compare performance across both datasets, we determined the set of variants which are found in both datasets, as well as in the genotype data of the Partners Biobank. We randomly selected 20, 000 loci with the *--thincount* option in PLINK2 (Purcell and Chang, 2020). Although *APOE* variants are known to have a very high effect size for AD, explaining around a quarter of the total heritability (Zhang et al., 2020), including the *APOE* region in a polygenic risk score or integrated risk model has been shown to be insufficient to account for the large risk attributed to *APOE* (Ware et al., 2020). To fine tune our integrated risk models on other *non-APOE* variants with much smaller effect sizes and good prediction power, we decided to keep *APOE* status as a separate predictor. At the same time, we made sure that the extended *APOE* region (from 45, 000, 000 − 46, 000, 000bp on chromosome 19) is excluded while the two *APOE* loci 19:45411941:T:C and 19:45412079:C:T are kept in the data. This leaves 18, 038 loci.

The final data used for the computation of the integrated risk models consists of these 18, 038 loci, as well as the following epidemiological covariates: age, sex, and *APOE* status with classes “none” (encoded as 0), “single e4” (encoded as 1), or “e4/e4” (encoded as 2).

In the following experiments, we considered the datasets of Kunkle et al. (2019) and Jansen et al. (2019) separately and extracted SNP weights based on corresponding effect sizes. Next, we withhold a proportion *p* ∈ {0.1, …, 0.9} of the pool of Partners genotyped subjects as a validation dataset to fit an integrated risk model with the aforementioned methods, or to tune the hyperparameters of the neural network. Finally, we evaluated their performance on the unseen proportion of the data (1 − *p*). We report the mean of absolute residuals 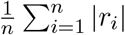 (where *n* is the number of subjects in the validation set and *r*_*i*_ is the residual for subject *i*), the AUC (Area under the ROC Curve), and the correlation between predicted and true outcomes.

Figure 1 shows results for the dataset of Kunkle et al. (2019). A series of observations are noteworthy. First, the mean of absolute residuals decreases with an increasing proportion of the data used for training, as expected.

**Figure 1:**
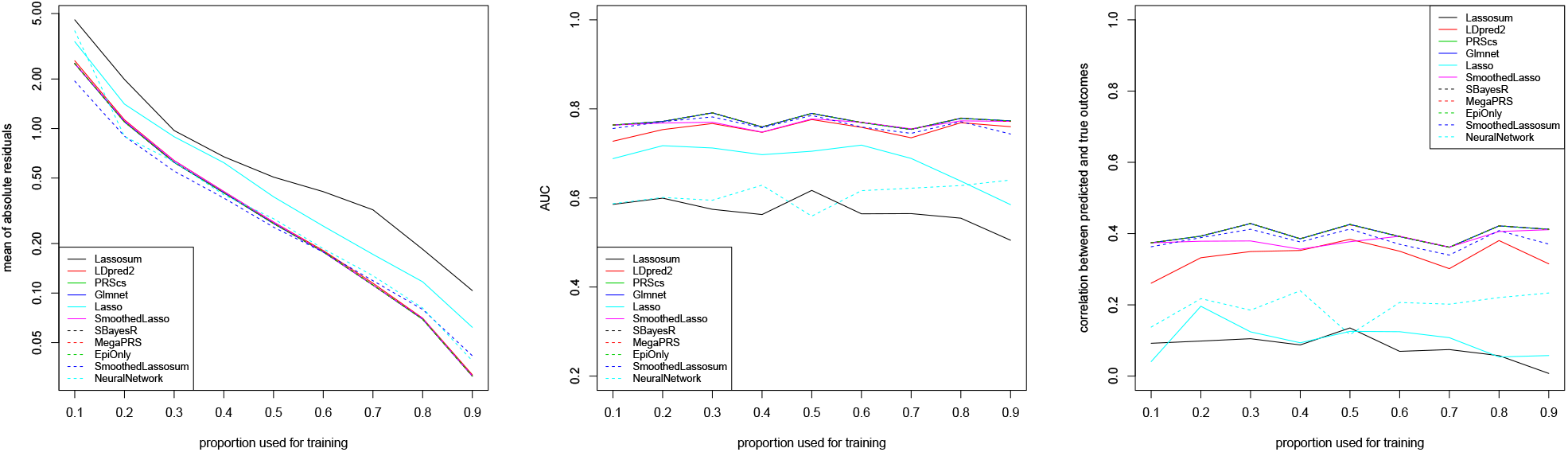
Dataset of clinically defined AD cases of Kunkle et al. (2019). Mean of absolute residuals (left), AUC (middle), and correlation between predicted and true outcomes (right) as a function of the proportion of data used for training. The behavior of most methods is similar to the one of LDpred2 or Glmnet.

Second, the AUC is very high (reaching almost 0.80) for all methods apart from Lassosum, Lasso, and NeuralNetwork. Interestingly, it is much less affected than the residuals by the proportion of data used for training and stays essentially constant for all training proportions. A similar picture is observed when looking at the correlation between predicted and true outcomes, which is roughly equally high for all methods apart from Lassosum, Lasso, and NeuralNetwork. After training, NeuralNetwork achieves a very low mean of absolute residuals, though its AUC and its correlation between predicted and true outcomes somewhat lacks behind the other methods. NeuralNetwork does manage to achieve an increased performance for higher proportions of training data (in both the AUC metric and with respect to the correlation between predicted and true outcomes). This is sensible, as neural nets traditionally need large amounts of data to be trained on.

Third, using epidemiological covariates only in a simple linear regression fit seems to perform very well on this dataset. This seems to suggest that actually, the response is well explained by the genetic factor of *APOE* status as well as the other non-genetic factors (such as age), and that the remaining genetic information is rather negligible for prediction.

Fourth, our proposed SmoothedLassosum considerably improves upon Lassosum of Mak et al. (2017), now drawing equal with state-of-the-art methodology such as LDpred2 with respect to e.g. the AUC measure. Moreover, our proposed SmoothedLassosum achieves a considerably improved mean of absolute residuals compared to Lassosum, and a state-of-the-art correlation between predicted and true outcomes. The reason for the reduced performance of Lassosum is not fully understood. However, it is likely related to the fact that Lassosum is not designed to incorporate epidemiological covariates (see Section 4 for more details).

The results for the dataset of Jansen et al. (2019), reported in Figure 2, are almost identical to the ones for the dataset of Kunkle et al. (2019) in Figure 1. In particular, the Lassosum, Lasso, and NeuralNetwork algorithms generally have the weakest performance on this dataset, while the other methods perform equally well. Importantly, SmoothedLassosum considerably improves upon Lassosum by achieving a mean of absolute residuals, AUC, and correlation between predicted and true outcomes that is similar to the others methods.

**Figure 2:**
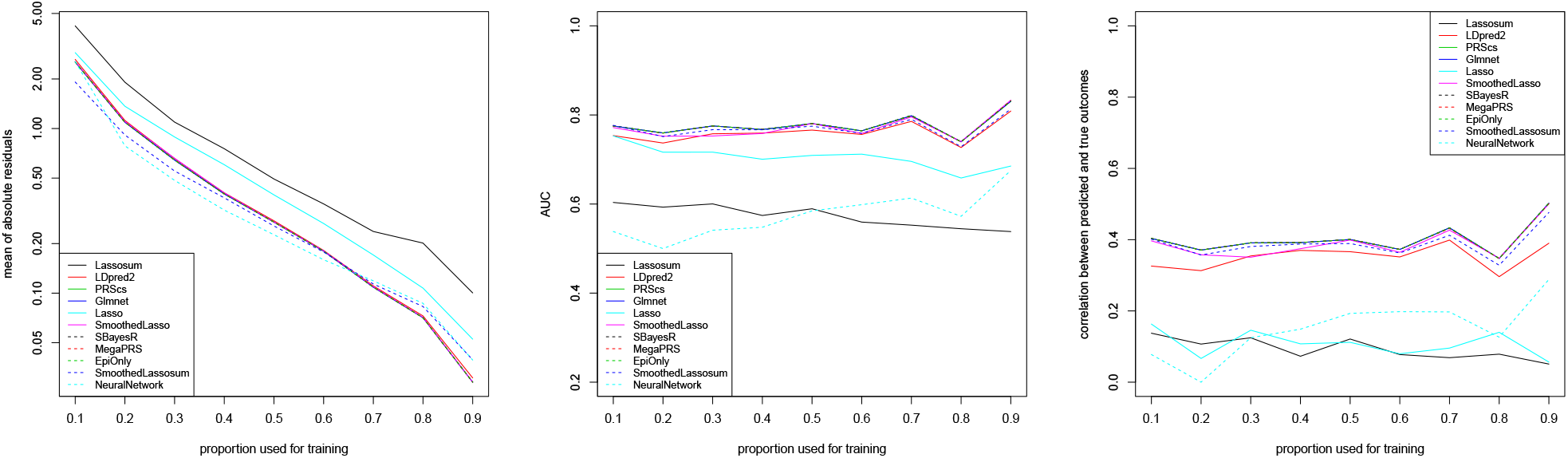
Dataset of AD-by-proxy phenotypes of Jansen et al. (2019). Mean of absolute residuals (left), AUC (middle), and correlation between predicted and true outcomes (right) as a function of the proportion of data used for training. The behavior of most methods is similar to the one of LDpred2 or Glmnet.

The very similar behavior of all methods is expected. The two experiments differ only in the way the response (AD status) is defined. The response provided in Kunkle et al. (2019) consists of clinically defined AD cases, while the one of Jansen et al. (2019) contains AD-by-proxy phenotypes which are based on 13 independent GWS loci having a strong genetic correlation of (at least) 0.81 with the AD status.

### 3.2 COPD study

The dataset considered in Section 3.1 is characterized through binary outcomes. In this section, we consider a continuous response in the context of Chronic Obstructive Pulmonary Disease (COPD). To be precise, we look at the COPDGene study of Regan E.A. (2010), a case-control study of COPD in current and former smokers (Silverman et al., 1998, 2000) which has been sequenced as part of the TOPMED Project. The data we employed are available through dbGaP (NHLBI TOPMed, 2018).

Chronic obstructive pulmonary disease (COPD) is the third leading cause of death in the United States (NHLBI TOPMed, 2018). The dataset we consider contains subjects with severe COPD, defined as having a *FEV1* ratio of *<* 40% predicted at an early age (*<* 53 years) without alpha-1 antitrypsin deficiency (a known Mendelian risk factor for COPD). The *FEV1* ratio, also called the Tiffeneau-Pinelli index describes the proportion of lung volume that a person can exhale within the first second of a forced expiration in a spirometry (pulmonary function) test. We focus on chromosome 15 and consider the risk loci for spirometric measures which have been identified in Lutz et al. (2015). Overall, we consider 8, 881 loci for 3, 495 individuals.

The genetic information is then matched to four epidemiological covariates. The final data used for the computation of the integrated risk models consists of the 8, 881 loci, as well as age, sex, pack-years of smoking, and height in centimeters. We aim to predict FEV1 from this data, again using a classic training (proportion *p* ∈ (0, 1)) and validation (proportion 1 − *p*) setup. We apply all algorithms as outlined in Section 3. As the AUC is only defined for a categorical response, we only report the mean of absolute residuals and the correlation between predicted and true outcomes.

Results of this experiment are given in Figure 3. We observe that measurements are overall more unstable than in Section 3.1, though as usual, the mean of absolute residuals in Figure 3 (left) decreases with an increasing proportion of the data used for training.

**Figure 3:**
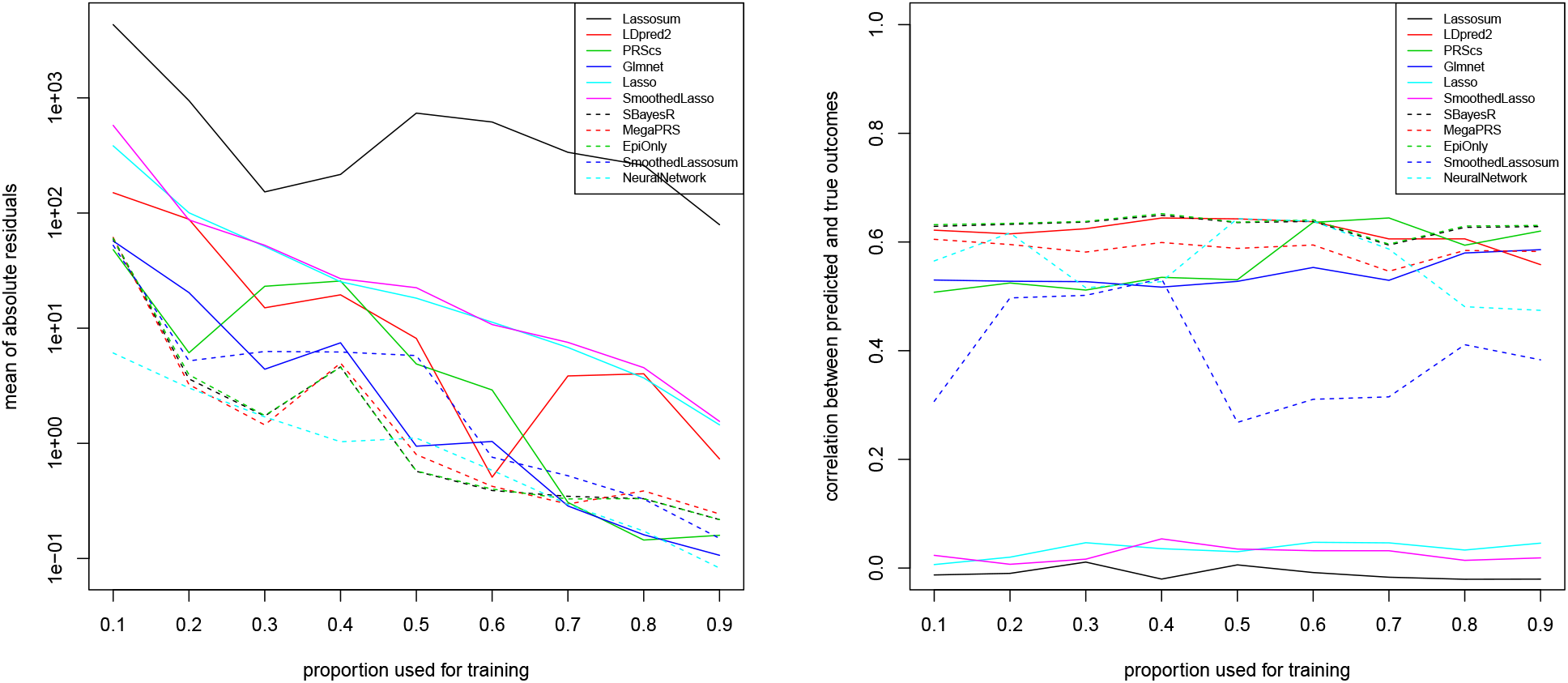
Dataset of AD-by-proxy phenotypes of Jansen et al. (2019). Mean of absolute residuals (left) and correlation between predicted and true outcomes (right) as a function of the proportion of data used for training.

Lassosum is again not performing at its best, which is likely related to the fact that we are aiming to predict a continuous response (see Section 4 for more details). The Lasso and SmoothedLasso approaches are performing average. Together with LDpred2 and PRScs, our proposed Smoothed-Lassosum approach performs very well and again considerably improves upon the original Lassosum. Glmnet is again one of the best methods together with SBayesR, MegaPRS, though a fit of epidemiological covariates only also seems to have high predictive power. NeuralNetwork seems to be very suited in this experiment to learn the continuous FEV1 responses from the input data.

The correlation between predicted and true outcomes, shown in Figure 3 (right), confirms that most state-of-the-art algorithms achieve a comparable correlation of around 0.6. The performance of our SmoothedLassosum is slightly worse than those methods with regards to the correlation between predicted and true outcomes, though it again considerably improves upon Lassosum (as well as Lasso and SmoothedLasso) which seem to have difficulties to predict the continuous FEV1 response from this data.

## 4 Discussion

This article considered the calculation of an integrated risk model by minimizing a smoothed version of the Lassosum objective function (see eq. (1)) introduced in Mak et al. (2017). Utilizing a smoothing approach circumvents the non-differentiability of the *L*_1_ penalty of Lassosum, thus allowing for an efficient minimization with quasi-Newton algorithms.

An experimental study on Alzheimer’s disease and COPD demonstrates that our smoothed Lassosum improves upon the original Lassosum of Mak et al. (2017), measured with respect to the mean of absolute residuals, the AUC, and the correlation between predicted and true outcomes, thus making it draw equal in accuracy with state-of-the-art approaches. The reduced performance of Lassosum we observe in the simulations is likely attributed to the fact that (a) Lassosum is not designed to incorporate epidemiological covariates in integrated risk models, and (b) Lassosum is not designed for continuous responses (as in the COPD study). In particular, although recomputing the SNP-wise correlation ***r*** = *X*^T^ ***y*** in eq. (1) and using them in place of ***y*** is a valid approach and an admissible input to the Lassosum objective function, the distribution of residuals is different from the one of the original binary response (without regressing out the covariates), which might cause a suboptimal behavior of the Lassosum algorithm. In contrast, our smoothed Lassosum works well in those cases.

Using an *L*_1_ penalty in eq. (1) has the advantage that, in analogy to the original Lasso of Tibshirani (1996), computing arg min_*β*∈ℝ*p*_ *L*(***β***) performs both regression of the polygenic risk score or integrated risk model and variable selection simultaneously. One potential drawback of our proposed smoothed Lassosum is that it yields dense minimizers (i.e., unused predictors are not necessarily shrunk to zero), meaning that the variable selection property is not preserved. This is not necessarily a disadvantage, as usually the fitted models are only used for risk prediction, for which our dense models achieve a high accuracy. Moreover, other widespread methods such as neural networks likewise do not provide variable selection. If necessary, sparseness can be restored after estimation via thresholding, meaning that all entries *β*_*i*_ of the estimate ***β*** of eq. (1) satisfying |*β*_*i*_| *< τ* for some threshold *τ* are set to zero. Determining an optimal threshold remains for future research.

## Acknowledgements

This work involved the use of the Enterprise Research Infrastructure & Services (ERIS) at Massachusetts General Hospital. We thank the MGB/Partners HealthCare Biobank for providing genomic and health information data.

Molecular data for the Trans-Omics in Precision Medicine (TOPMed) program was supported by the National Heart, Lung and Blood Institute (NHLBI). See the TOPMed Omics Support Table (Table 1) for study specific omics support information. Core support including centralized genomic read mapping and genotype calling, along with variant quality metrics and filtering were provided by the TOPMed Informatics Research Center (3R01HL-117626-02S1; contract HHSN268201800002I). Core support including phenotype harmonization, data management, sample-identity QC, and general program coordination were provided by the TOPMed Data Coordinating Center (R01HL-120393; U01HL-120393; contract HHSN268201800001I). We gratefully acknowledge the studies and participants who provided biological samples and data for TOPMed.

**Table 1:** TOPMed Omics Support Table. Broad Genomics = Broad Institute Genomics Platform; Broad Metabolomics = Broad Institute and Beth Israel Metabolomics Platform; Keck MGC = Keck Molecular Genomics Core Facility; NWGC = Northwest Genomics Center.

| TOPMed<br>Accession # | TOPMed<br>Project | Parent Study<br>Short Name | TOPMed<br>Phase | Omics Center | Omics Support | Omics Type |
| --- | --- | --- | --- | --- | --- | --- |
| phs000951 | COPD | COPDGene | 1 | NWGC | 3R01HL089856-08S1 | WGS |
| phs000951 | COPD | COPDGene | 2 | Broad Genomics | HHSN268201500014C | WGS |
| phs000951 | COPD | COPDGene | 2.5 | Broad Genomics | HHSN268201500014C | WGS |
| phs000951 | COPD | COPDGene | 4 | NWGC | HHSN268201600032I | RNASeq |
| phs000951 | COPD | COPDGene | 5 | NWGC | HHSN268201600032I | Methylomics |

Parent Study-specific Acknowledgements:

- NHLBI TOPMed: Childhood Asthma Management Program (CAMP)
- NHLBI TOPMed: Genetic Epidemiology of COPD Study (COPDGene). The COPDGene project described was supported by Award Number U01 HL089897 and Award Number U01 HL089856 from the National Heart, Lung, and Blood Institute. The content is solely the responsibility of the authors and does not necessarily represent the official views of the National Heart, Lung, and Blood Institute or the National Institutes of Health. The COPDGene project is also supported by the COPD Foundation through contributions made to an Industry Advisory Board comprised of AstraZeneca, Boehringer Ingelheim, GlaxoSmithKline, Novartis, Pfizer, Siemens and Sunovion. A full listing of COPDGene investigators can be found at: http://www.copdgene.org/directory

## Data Availability Statement

The genotype data used in the simulations is available from the Partners Biobank (Partners, 2020). The summary statistics of Kunkle et al. (2019) and Jansen et al. (2019) used in the simulations are available online, see NIAGADS (2016) and CTG Lab (2021).

The data that support the findings of this study are openly available in “NHLBI TOPMed: Boston Early-Onset COPD Study in the National Heart, Lung, and Blood Institute (NHLBI) Trans-Omics for Precision Medicine (TOPMed) Program” at https://www.ncbi.nlm.nih.gov/projects/gap/cgi-bin/study.cgi?study_id=phs000946.v3.p1.

## A Principal component plots

Figures 4 and 5 show the first eight principal components of the HRC-imputed genotype data downloaded from Partners Biobank. All individuals we kept in the dataset are self-reported nonhispanic white (NHW) individuals. We excluded outliers which are at least 5 standard deviations away from the mean value of each of the ten principal components. In Figure 4 we observe a negligible amount of stratification based on the genotyping chip, but given the even distribution of cases/controls across chips displayed in Figure 5, this should not affect the results.

**Figure 4:**
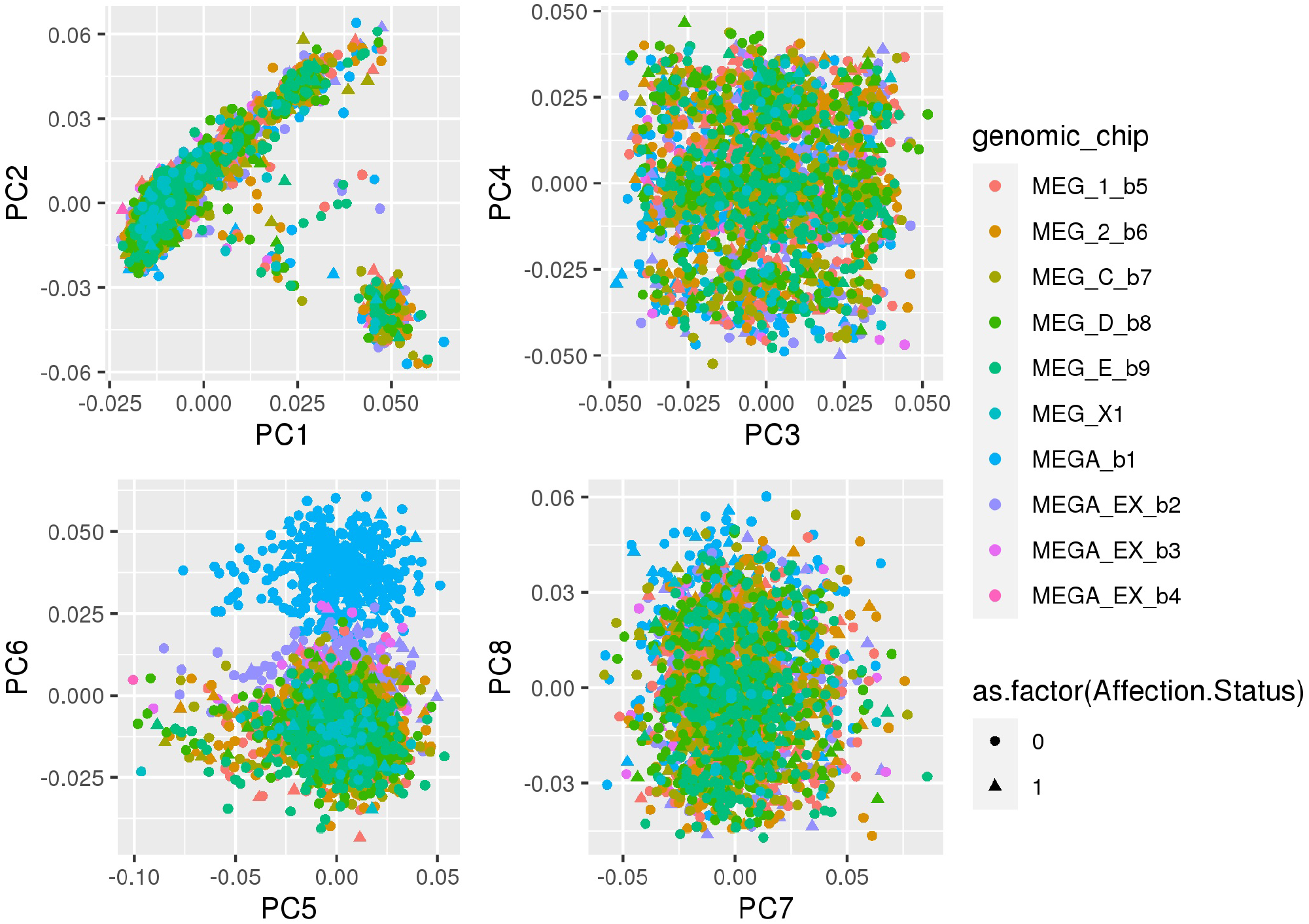
First eight principal components of the HRC-imputed genotype data downloaded from Partners Biobank. Stratification by genomic chip and affection status.

**Figure 5:**
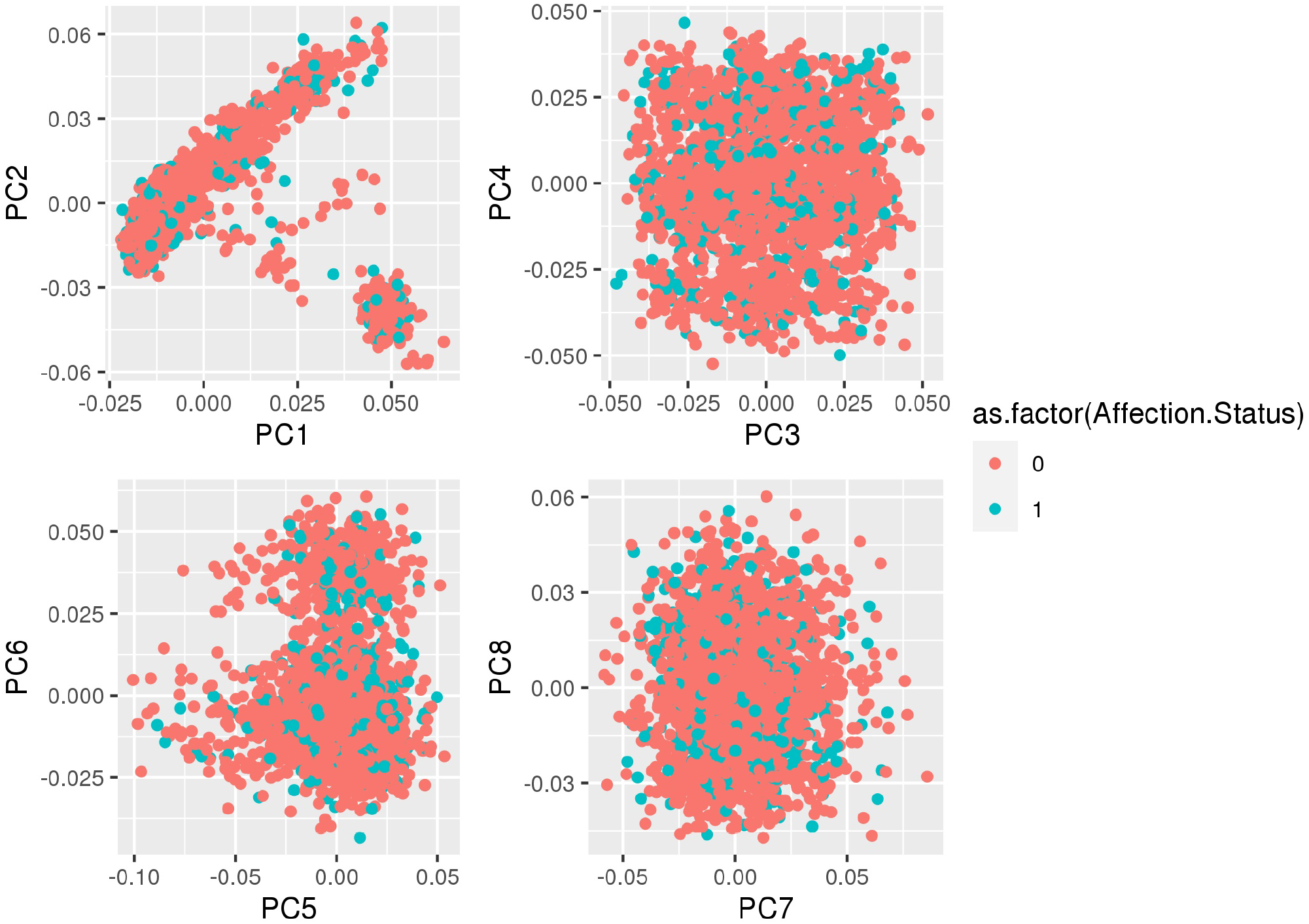
First eight principal components of the HRC-imputed genotype data downloaded from Partners Biobank. Stratification by affection status.

